# Avoidance of simultaneous patch use in Japanese large-footed bats

**DOI:** 10.64898/2026.02.09.704905

**Authors:** Emyo Fujioka, Masashi Shiraishi, Tamao Hirao, Yui Onishi, Dai Fukui, Shizuko Hiryu

## Abstract

Group foraging can enhance prey detection, but depending on resource availability, it may also generate conflicts among conspecifics. To understand how animals balance these benefits and costs, foraging performance must be evaluated together with inter-individual interactions. However, under fully natural conditions, it remains challenging to quantify both simultaneously. Here, we investigated how individual foraging efficiency and pairwise interactions are shaped when more than one individuals simultaneously exploit the same foraging patch, using the Japanese large-footed bat (*Myotis macrodactylus*) as a model system. We monitored an entire pond functioning as a natural foraging patch using two thermal cameras and an eight-channel microphone array, and reconstructed the arrival, prey-attack, and exit times of individual bats. Using a Poisson generalized linear mixed model (GLMM), we found that prey-attack rates were approximately 25% lower during paired flights than during solitary flights. We then constructed a null model in which arrival, attack, and departure events followed independent Poisson processes parameterized from the empirical data. Compared with null-model predictions, both the total duration and the duration of individual paired flights in the empirical data were significantly shorter, indicating that bats limited the time spent co-using the same patch relative to solitary foraging. In addition, the probability that the first exiting individual was the one that arrived earlier or later did not deviate from chance levels, providing no evidence for a prior residence advantage. Together, these results demonstrate that simultaneous patch use avoidance occurs independently of arrival order and coincides with reduced prey-attack rate, suggesting that bats leave shared patches and move to alternative foraging sites to mitigate losses in prey-attack efficiency. Our findings highlight bats as an excellent model system for non-invasively linking individual behavior and foraging performance via echolocation, and for elucidating the dynamics of foraging behavior and sensory interference in the wild.

## Introduction

Foraging behavior is one of the most fundamental activities in animals and directly affects survival and fitness [1, 2]. Many predatory species do not forage exclusively alone but instead adopt strategies involving temporary or permanent group formation (e.g., fish[3, 4]; ants[5, 6]; birds[7, 8]). Such strategies confer multiple advantages, including an expansion of individual perceptual ranges that enhances prey-detection capabilities [9, 10], improved access to temporally and spatially ephemeral food patches [11, 12], and increased foraging success through information transfer among group members, which facilitates efficient food searching and hunting [13, 14]. At the same time, group foraging entails costs, such as intensified intraspecific competition and interference arising from resource sharing [15, 16], as well as reductions in per capita food intake [17, 18]. Although these trade-offs have been demonstrated in numerous studies at the individual level (e.g., [10, 12]) and under controlled conditions (e.g., [14, 17]), quantitatively observing the foraging behavior and interactions of multiple freely moving individuals in fully natural environments, and understanding how predation is actually carried out under such conditions, remain extremely challenging.

Bats rely on echolocation for navigation and prey detection in darkness [19, 20]. Echolocation calls contain rich information about the direction of attention and the behavioral state of an individual. For example, feeding buzzes emitted immediately prior to prey capture indicate when a bat is capturing target prey, and the interval of ultrasonic emissions allows precise identification of the timing at which prey-approach behavior is initiated. Thus, in bats, echolocation calls enable high-resolution quantification of foraging behavior under fully natural conditions [21, 22]. In particular, the Japanese large-footed bat (*Myotis macrodactylus*) uses echolocation to detect aquatic insects near the water surface and captures prey by trawling flight over ponds and streams [23, 24]. This behavioral specialization facilitates fixed-site observations of a foraging patch, making this species an excellent wild model for quantitatively analyzing foraging behavior and inter-individual interactions based on acoustic recordings.

A distinctive feature of inter-individual interactions during bat foraging is that echolocation calls and their echoes are audible not only to the emitting individual but also to nearby conspecifics. Listening to the calls of others can yield both benefits and costs during foraging. For example, it has been reported that bats—including both conspecifics and heterospecifics—eavesdrop on the echolocation calls of others to obtain profitable information about foraging sites [25-28]. At the same time, calls emitted by nearby individuals can interfere with a bat’s own echoes, potentially degrading echolocation performance and reducing foraging efficiency. Indeed, recent studies have highlighted a trade-off in group foraging, whereby foraging with a small number of conspecifics can facilitate prey detection, whereas the presence of too many individuals can instead reduce efficiency [29]. Considering these opposing benefits and costs, understanding bat foraging strategies when multiple individuals occupy the same foraging patch requires quantitative evaluation not only of individual behavior but also of its relationship with local prey resources. In Japanese large-footed bats, aquatic insects targeted during foraging emerge at the water surface and exit shortly thereafter, such that competition is expected among individuals attempting to exploit prey immediately after emergence. Accordingly, analyses of foraging behavior in this species provide an opportunity to reveal dynamic inter-individual interactions and strategic adjustments within a shared foraging patch.

In this study, we monitored the entire pond as a natural foraging site and examined the foraging behavior of Japanese large-footed bats that arrived sequentially at the site, with particular attention to periods in which more than one individual was co-using the site simultaneously. At the focal pond, both solitary foraging events and situations in which additional bats arrived and foraged simultaneously were observed. In echolocation-based foraging targeting prey at the water surface, acoustic interference arising from reflections off the water surface is expected to occur—an effect that is absent during foraging in featureless three-dimensional open space. Consequently, when the foraging patch is spatially restricted, sharing the patch with conspecifics is predicted to impose substantial echolocation-related costs on Japanese large-footed bats. Based on this reasoning, we hypothesized that the presence of other bats within the same foraging patch reduces foraging efficiency and that individuals adopt behavioral strategies to avoid co-using the patch with conspecifics.

To test this hypothesis, we deployed wide–field video cameras and a microphone array configured to cover the entire pond, allowing us to record the arrival and departure times of successive bats as well as their prey-capture behavior within the foraging patch. We further modeled bat arrival, prey-attack, and exit events at the patch as Poisson processes and estimated their occurrence rates from the empirical data to construct a null model that excludes inter-individual interactions. By comparing the empirical data with this null model, we evaluated whether bats adjust their behavior in response to the presence of conspecifics when co-using the same patch—that is, whether patch staying times deviate from expectations under a simple probabilistic process. The primary objective of this study was to investigate how Japanese large-footed bats adjust their behavior under natural conditions when more than one individual simultaneously exploits the same foraging patch and to clarify the relationship between such behavioral adjustments and foraging efficiency. To further address this objective, we additionally analyzed which individual was more likely to leave first during simultaneous multi-individual foraging bouts, providing complementary insights into inter-individual interactions during bat foraging.

## Materials and methods

### Target species and study site

Target species of this study is *Myotis macrodactylus*. This bat species performs echolocation using frequency-modulated calls [23, 30]. When attacking insect prey, individuals emit a feeding buzz, during which the call frequency decreases immediately prior to the attack [23, 24]. This indicates that the timing of prey-attack events can be identified from the recorded echolocation sounds. We defined foraging efficiency as the frequency of attack attempts on prey, that is, the encounter rate with prey.

The study site was a single pond (approximately 20 × 20 m) located within the Tomakomai Experimental Forest of Hokkaido University (Fig. 1), which is one of several ponds distributed along Horonai stream within the forest. At this pond, *M. macrodactylus* have been observed foraging just above the water surface alone or in the presence of conspecifics [24]. Field recordings were conducted on three nights (12, 16, and 17 September 2024). Data collection began shortly after sunset (approximately 17:30), when the first bat appeared, and continued for approximately one hour each night.

**Fig. 1.**
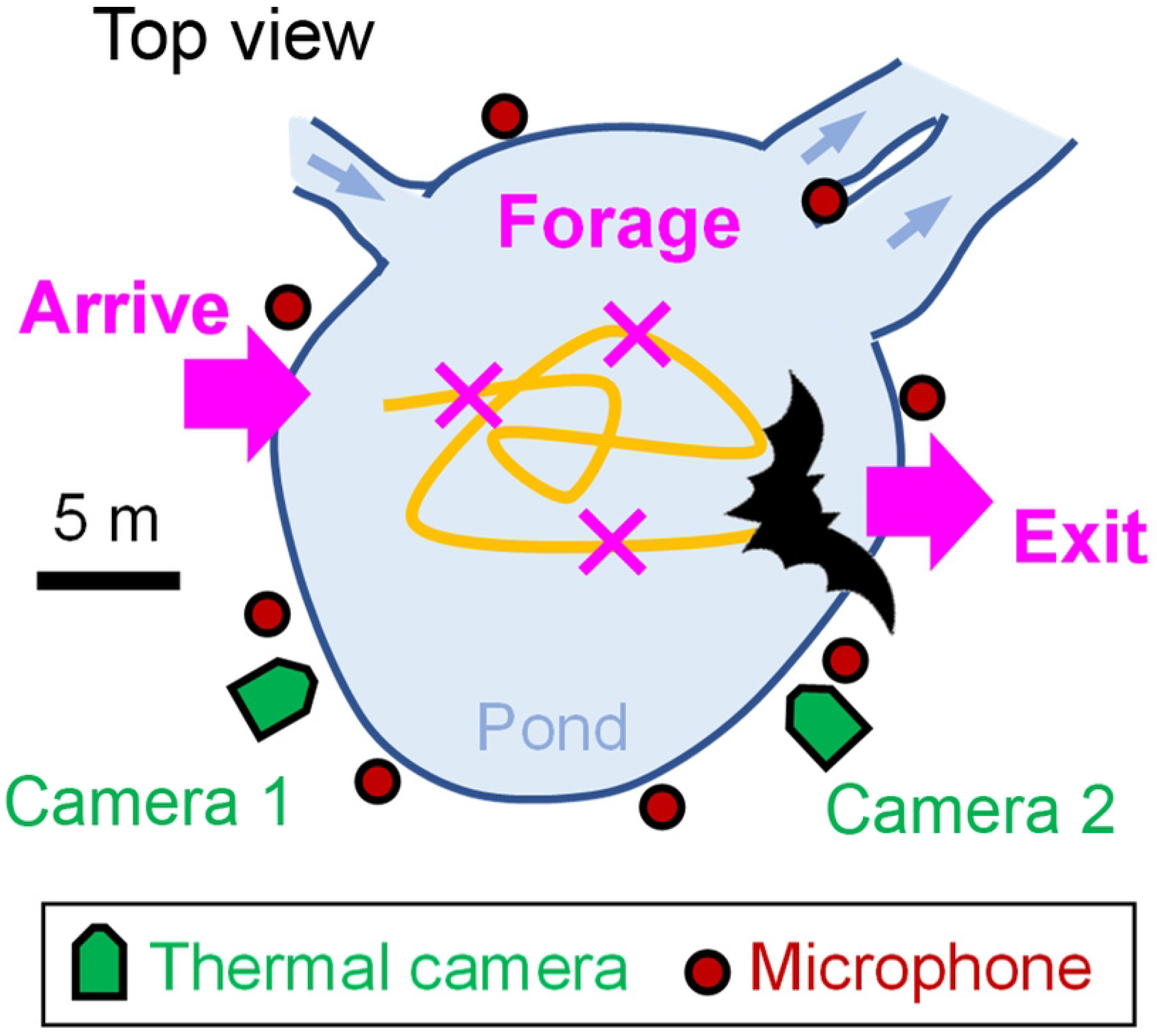
Schematic diagram of the study site and measurement system. Blue arrows show the direction of the stream.

Although the study site is designated as a wildlife reserve by the local government, no specific permits were required for this study because it was a non-invasive observational study that did not involve any endangered or protected species, animal capture, or habitat disturbance. Experiments at the pond were conducted with permission from the forest administration.

### Video and acoustic recording

For video recordings, two FLIR A65 thermal video cameras (field of view: 90°, frame rate: 30 fps; Teledyne FLIR LLC, Oregon, USA) were used to record bat arrivals at and exits from the pond as monochrome video files on a personal computer (Fig. 1). The recordings from the two cameras were temporally synchronized by clapping hands within the overlapping fields of view of both cameras. Using the recorded videos, the arrival and exit times (hh:mm:ss) of successive bats were extracted, and inter-arrival intervals and patch residence times in patch for each bat were calculated.

For acoustic recordings, eight ultrasonic-band MEMS microphones (custom-made, based on the SPU0410LR5H; Knowles) were deployed surrounding the pond (Fig. 1). This configuration enabled comprehensive recording of all vocalizations produced within the pond, including feeding buzzes emitted by individual bats. Acoustic signals captured by the microphones were recorded on a personal computer via a data acquisition device (USB-6356; National Instruments, Inc., USA) at a sampling rate of 500 kHz. Audio and video recordings were synchronized using the handclap sounds that were also used for video synchronization.

The timing of prey-attack events was determined from the acoustic data by identifying the terminal portion of feeding buzzes in which call frequency decreased, with a temporal resolution of 0.1 s. When no frequency decrease was observed, the event was recorded as a non-attack [24]. In this study, situations in which only a single bat was present within the foraging area were defined as the single-bat context, whereas situations involving two or more bats were defined as the multiple-bat context. In the multiple-bat context, the individual flying near the microphone that recorded the feeding buzz was determined to be the emitter of the feeding buzz. When it is hard to identify the feeding-buzz emitter using such snapshot of video and acoustic recordings because inter-individual distance was too small, the emitter was identified using time differences of arrival (TDOA) in the microphone array. Specifically, the relative spatial positions of the bats were determined from the temporal patterns of TDOA in acoustic sequences containing a feeding buzz recorded by the microphones near the bats. Then, the bat that emitted feeding buzz was identified by comparing the acoustically determined relative positions with the bats’ flight positions observed in the synchronized video recordings.

### Null model construction and statistical analysis

We constructed a stochastic behavioral simulation model in MATLAB (MathWorks, USA), in which three events at the foraging site—arrival, prey attack, and exit—were treated as independent Poisson processes. This model was positioned as a null model that does not incorporate inter-individual interactions, and the event rates (occurrence probabilities per unit time) for all processes were derived entirely from the empirical data obtained in this study. Specifically, the arrival rate was defined as the number of bats arriving at the foraging site per unit time. The prey-attack rate was estimated separately for the single-bat and multiple-bat contexts by modeling the number of prey attacks per unit time for each bat using a Poisson generalized linear mixed model (GLMM), with the contexts (i.e., single/multiple) treated as a fixed effect (see Section 2.4 for details). The exit rate was defined as the inverse of the mean seconds of patch residence time of bats at the foraging site.

Because no significant day-to-day differences were detected in inter-arrival intervals or the patch residence time (all *p* > 0.05, Kolmogorov–Smirnov tests), these rates were estimated using pooled data from all three days. In contrast, inter-attack intervals differed significantly among days (*p* < 0.001, Kolmogorov–Smirnov test), and therefore day-specific attack rates were used in the model.

Differences in prey-attack rate between the single-bat and multiple-bat contexts were analyzed using a Poisson GLMM, in which the number of prey attacks per individual was treated as the response variable and patch residence time of individuals was included as an offset term. Context (single-bat vs. multiple-bat) was specified as a fixed effect, and the significance of its effect was evaluated using a Wald test. For individuals that engaged in multiple-bat context, attack events were assigned separately to the single-bat and multiple-bat contexts when the same individual flew alone before or after multiple-bat context. Individual identity and experimental date were included as random effects in the model.

Simulations of the mathematical null model were conducted with each trial covering the same duration as the empirical measurements (i.e., 1 h). A total of 10,000 trials were performed to generate the distributions of flight patch residence time under the single-bat and multiple-bat contexts, which were then tested for differences from the empirical data. Using the outputs of the mathematical model as the null distribution, deviations of the empirical bat data from model expectations were evaluated using a parametric bootstrap test.

To examine whether individuals tended to occupy the foraging patch in the multiple-bat context, we analyzed which bat exited first during paired flights: the individual that arrived earlier or the one that arrived later. When multiple bats arrived at or exited from the foraging patch nearly simultaneously (with a time difference of less than 1 s), the order of events could not be reliably determined; such cases were therefore excluded from this analysis. To restrict the analysis to simple and interpretable situations, only two-bat conditions (i.e., paired flights) were considered. Individuals that experienced situations involving three or more bats were excluded from this analysis (observed only in the case of numerical simulation, see Results).

## Results

Over the three observation nights (12, 16, and 17 September), the first individual appeared at approximately 18:30, and a total of 44, 75, and 61 individuals were recorded within the subsequent 1 h, respectively (Fig. 2A, S1 Dataset). Bats arrived at the foraging patch successively, repeatedly attacked prey, and then exited. The inter-arrival interval, the patch residence time, and inter-attack interval averaged 57.6 ± 46.0 s, 17.3 ± 17.9 s, and 2.52 ± 2.14 s (mean ± SD), respectively, across the three days (Fig. 2B). A total of 31 flight events of multiple-bat context were observed over the three days, and no cases involving three or more individuals were recorded.

**Fig. 2.**
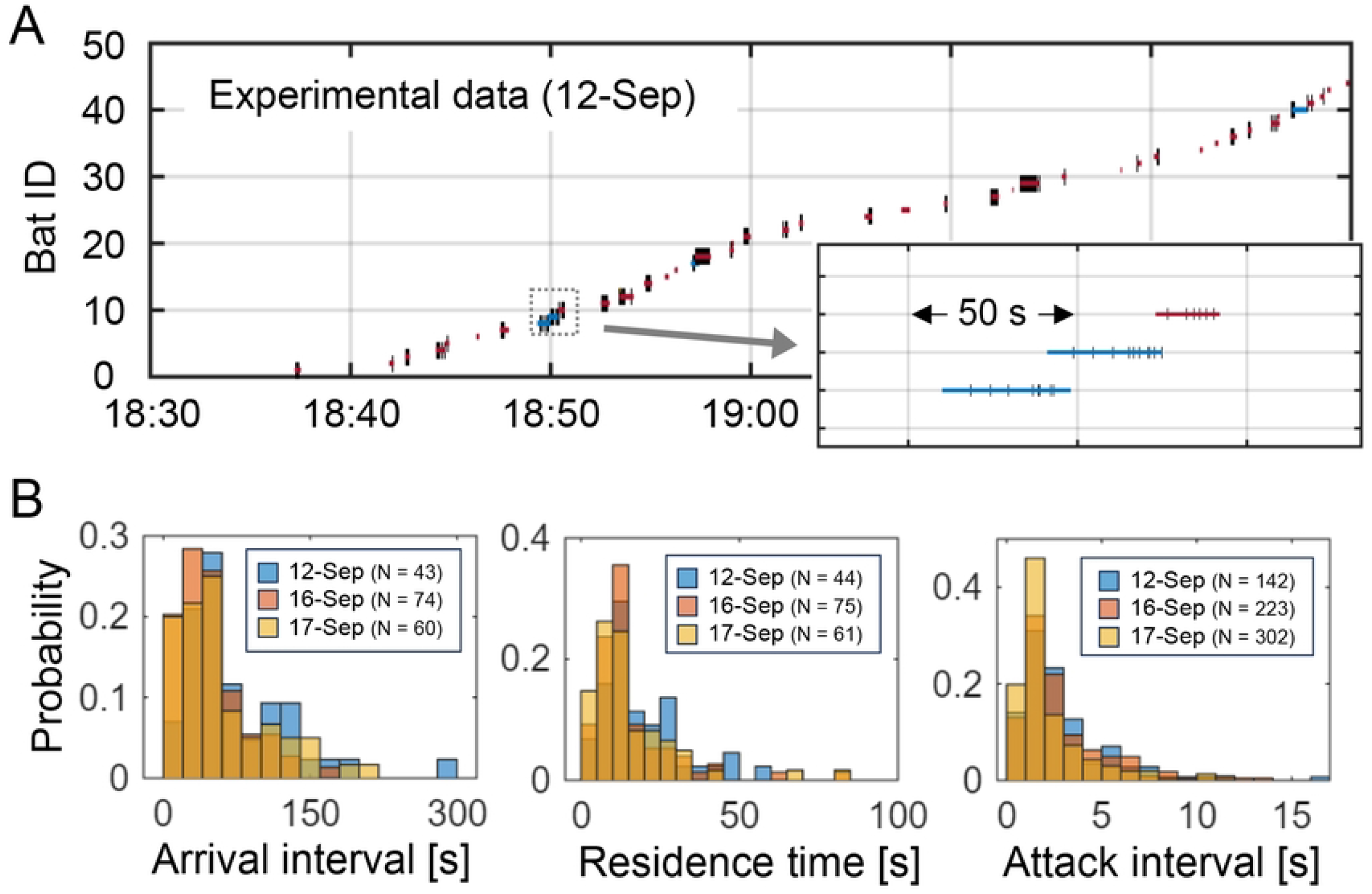
(A) Time series data of the bats foraging at the pond during 1-h measurement. The length of the line indicates the time spent at the pond. Red lines indicate the bats exiting the pond alone, whereas the blue and yellow lines show the bats exiting the pond in group; the latter bats entered before (blue) and after (yellow) the entrance of the former bats. The vertical lines across the horizontal lines indicate the timing of attacking prey. (B) Proportional distribution of arrival intervals (left), patch residence time (middle) and attack intervals (right) of the bats during the measurement periods for each day.

The Poisson GLMM of prey-attack counts indicated that bats attacked prey at a mean rate of 0.224 attacks s^−1^ in the single-bat context (estimate = −1.498 ± 0.113 SE, z = −13.21, *p* < 0.001), and that this rate was significantly reduced in the multiple-bat context (estimate = −0.285 ± 0.131 SE, z = −2.17, *p* < 0.05; Fig. 3). Consequently, the prey-attack rate in the multiple-bat context was reduced by 25% relative to the single-bat context (exp[−0.285] = 0.75; S2 Table).

**Fig. 3.**
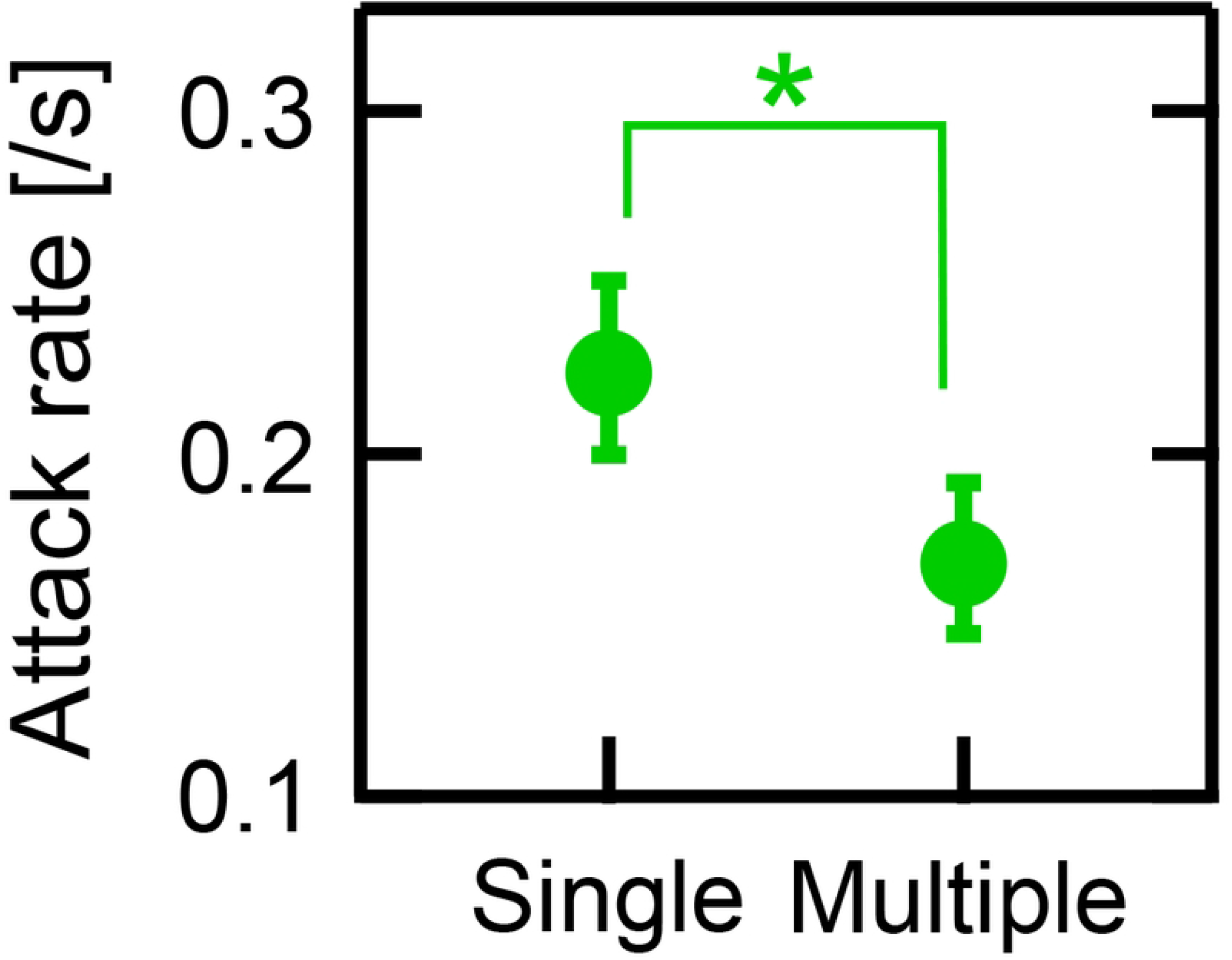
Prey attack rates during single and multiple flights estimated by a GLMM; The vertical lines show the standard errors. The slope of the fixed effect for multiple vs. single was -0.285 (p < 0.05 (^*^), Wald test). Number of data was 217 (single: 160, group: 57).

Using the parameters estimated from the empirical data, we conducted foraging simulations (Fig. 4A). The simulated outputs—inter-arrival intervals, patch residence time, and prey-attack rates per simulation run (i.e., 1 h)—closely matched the corresponding empirical values used as model inputs (S3 Table). In the multiple-bat context, such events occurred 13.1 ± 4.0 times per hour (mean ± SD, N = 10,000 simulations), with a total duration of 130.5 ± 52.6 s per hour. In contrast, the total duration of the multiple-bat context in the empirical data was 49 s (Day 1, N = 6), 79 s (Day 2, N = 13), and 61 s (Day 3, N = 12), which was significantly shorter than predicted by the simulations (Z ≈ −2.35, p < 0.001, one-sample Z test; Fig. 4B). Furthermore, the duration of individual multiple-bat contexts averaged 9.9 ± 10.0 s in the simulations (N = 131,374 events from 10,000 simulations), whereas the corresponding values in the empirical data were 5.5 ± 3.7 s (Day 1, N = 6), 6.2 ± 6.9 s (Day 2, N = 13), and 5.1 ± 3.0 s (Day 3, N = 12; mean ± SD). Parametric bootstrap tests revealed that the durations of individual multiple-bat contexts in the empirical data were significantly shorter than those predicted by the model for all days (p < 0.001; Fig. 4C).

**Fig. 4.**
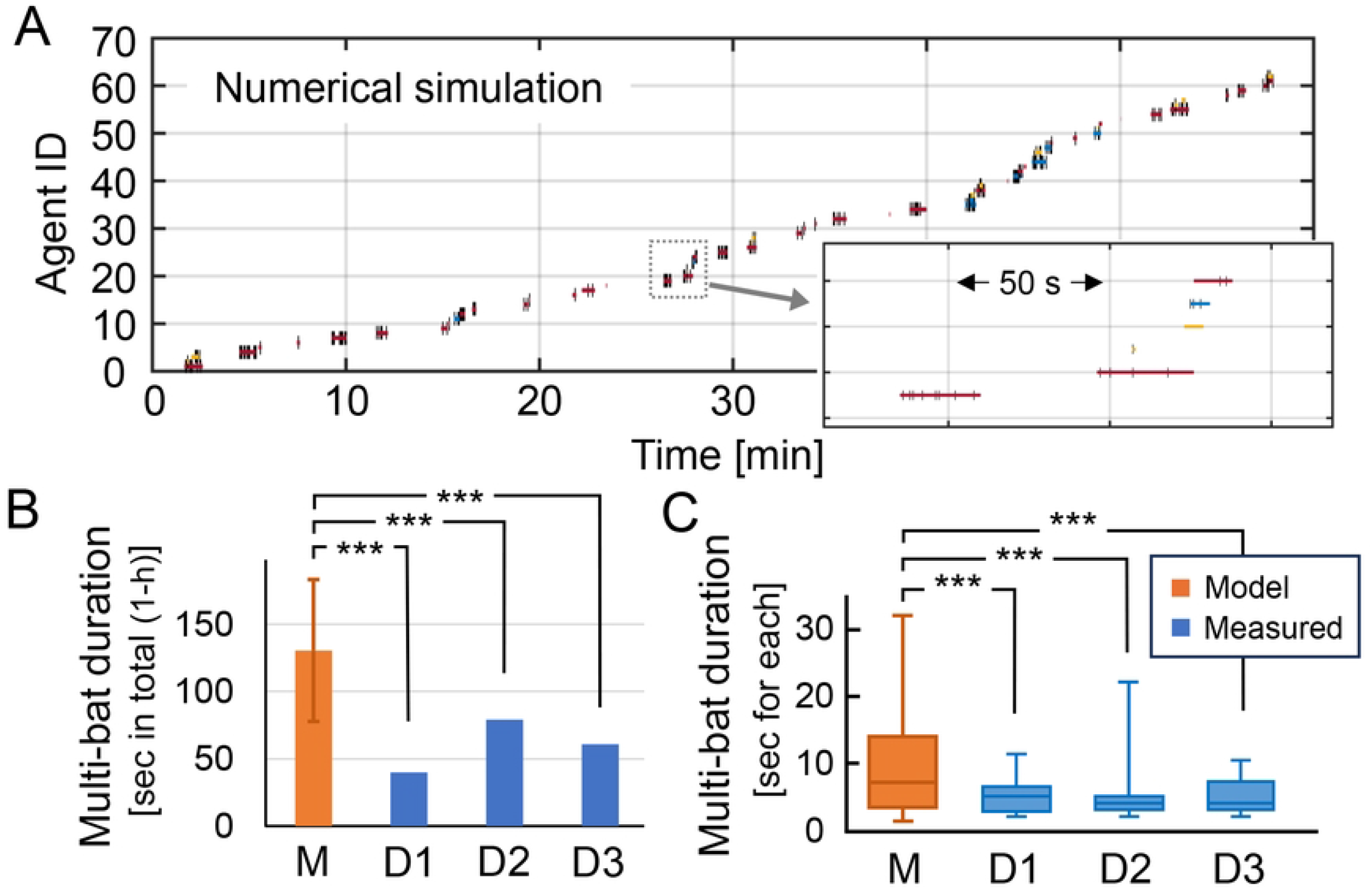
(A) Time series data of the 1-h foraging simulation of the null model (drawing rule is same as Fig. 2A). (B) Total time spent in the foraging patch of model and measured (three different days) bats while in multiple contexts. Vertical lines on the orange bar indicate the standard deviation based on 10,000 numerical simulations. One-sample Z test was conducted; ^***^ p < 0.001. (C) Boxplots (0.025, 0.25, 0.5, 0.75, and 0.975 quantiles) showing the consecutive durations of each multiple flight obtained from the numerical simulations (orange) and from three days of field observations of bats (blue). Parametric bootstrap tests were conducted; ^***^ p < 0.001.

In paired flights, we examined whether the individual that exited first was the one that arrived earlier (Bat A) or the one that arrived later (Bat B). In the simulations, once a paired flight began, both bats had identical departure probabilities, resulting in an equal likelihood of Bat A or Bat B exiting first (Fig. 5, left). Analysis of the empirical data revealed an exactly even split, with the first exit occurring equally often for Bat A and Bat B (Fig. 5, right; N = 20; Bat A = 10 cases, Bat B = 10 cases).

**Fig. 5.**
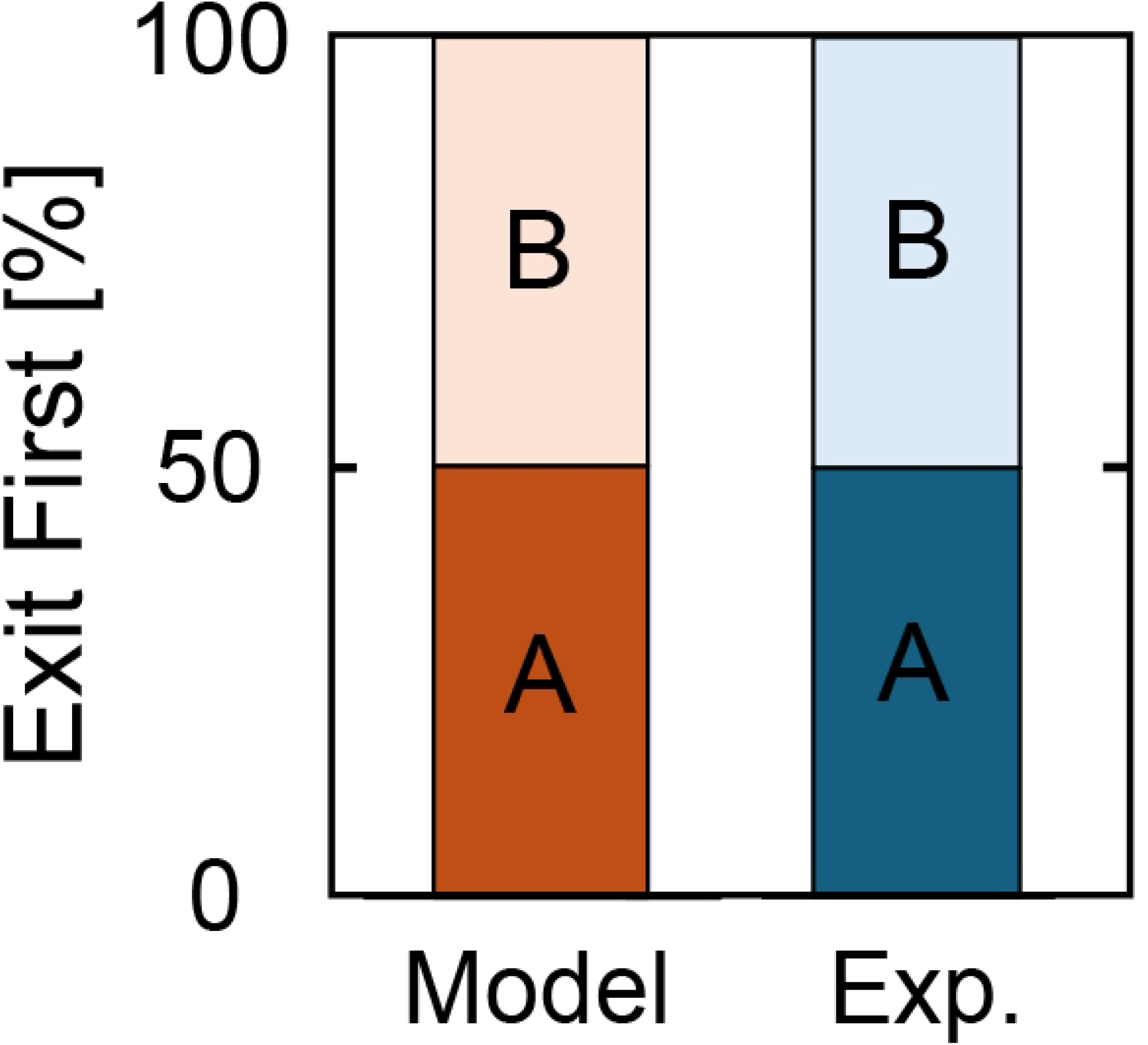
The bat that exited the pond first when in paired flights for the case of model simulation (red) and the experimental data (blue). Bat A is the bat that was staying before the paired flight together with the latter bat (Bat B). The numerical simulation was exhibited 10,000 times.

## Discussion

In this study, we used a microphone array and video cameras to quantify the sequential arrival, foraging, and exiting of bats within a foraging patch. Our results demonstrate that when two individuals were present simultaneously within the patch, prey-attack rate was reduced relative to single-bat flight context (Fig. 3). Because the pond we measured was sufficiently small that flight areas overlapped during paired flights, this reduction is likely attributable to the presence of conspecifics acting as moving obstacles and/or to acoustic interference between the ultrasonic signals emitted for echolocation, which may have decreased prey-detection efficiency. Furthermore, comparison with the null model we constructed revealed that the duration of paired flights was significantly shorter in the empirical data than predicted by the model (Fig. 4). This finding indicates that bats actively exited from the foraging patch to avoid simultaneous foraging with conspecifics. The study site was located within a university experimental forest, where multiple ponds and stream pools are distributed in the surrounding area. It is therefore plausible that bats avoided simultaneous foraging with conspecifics at a single patch and instead moved to alternative nearby foraging patches.

When two individuals are co-using a foraging patch, an important question is which individual leaves first in order to avoid simultaneous foraging. Previous studies have frequently reported behaviors in which individuals expel others from feeding sites (food patch defense), with the resident typically displacing the intruder—a pattern known as the prior residence effect [31, 32]. In the present study, however, the probability that the first individual to exit was the resident or the intruder did not differ from chance levels (Fig. 5), indicating that avoidance of simultaneous patch use does not depend on the order of arrival at the foraging patch. Accordingly, a prior residence effect does not appear to operate in the foraging behavior of Japanese large-footed bats. More generally, avoidance of simultaneous foraging with conspecifics at a feeding patch may be influenced not only by prior residence effects but also by kin selection [33] and dominance hierarchies [34]. High relatedness can promote tolerance or food sharing among individuals, whereas dominance relationships may lead to displacement or exclusion from feeding sites [35, 36]. To gain a deeper understanding of the mechanisms underlying collective foraging in bats, future studies should jointly examine foraging behavior, social hierarchy, and relatedness within colonies.

Japanese large-footed bats have been reported to emit various types of social calls at foraging sites [37], some of which are suggested to function in repelling conspecifics [38]. Thus, social calls could potentially influence the patch residence time and prey-attack rate during paired flights in the present study. However, social calls were observed infrequently, occurring in only 5 of the 31 paired flights (16.1%). When we reanalyzed the data using a Poisson GLMM after excluding flights in which social calls were observed (including those individuals during solitary flights), bats attacked prey at a mean rate of 0.225 attacks/s during solitary flights (estimate = −1.491 ± 0.112 SE, z = −13.36, *p* < 0.001), and this rate was significantly reduced during paired flights (estimate = −0.332 ± 0.147 SE, z = −2.26, *p* < 0.05). Consequently, prey-attack rate during paired flights was reduced by 28% relative to solitary flights (exp[−0.332] = 0.72; S2 Table). These results indicate that, regardless of the presence or absence of social calls, prey-attack rate was significantly lower during paired flights than during solitary flights, and the magnitude of this reduction was largely unchanged. This suggests that social calls likely had little effect on foraging efficiency in the present study.

Foraging efficiency during multiple-bat flights with conspecifics is likely to depend on the size of the foraging patch. If bats can fly within a patch while experiencing minimal interference from other individuals—such as overlap in flight paths or sensory interference—then the presence of conspecifics may not necessarily reduce prey-attack rate. Even in large patches, however, increasing the number of individuals can intensify competition, prompting predators to move to alternative foraging patches; as a result, the benefits gained at each patch tend to become approximately equal among individuals. Such distributions of predators across feeding sites are described by the ideal free distribution [39]. In bat research, it has been reported that increases in bat density at a given foraging patch lead to competition and subsequent shifts to alternative patches [40]. In the present study, because multiple ponds are distributed along the river within the experimental forest where our study pond is located, Japanese large-footed bats may similarly compare prey availability between the focal pond and other nearby foraging patches and avoid paired flights in order to maintain foraging efficiency.

In conclusion, our study demonstrates that prey-attack rate of *M. macrodactylus* declines during paired flights within a foraging patch and that bats actively avoid simultaneous foraging with conspecifics within the patch. By quantitatively characterizing foraging behavior that is otherwise difficult to capture under natural conditions, this study highlights the Japanese large-footed bat as an excellent model system for elucidating the dynamics of collective foraging and the mechanisms of sensory interference in the wild. These findings provide a foundation for future studies incorporating multiple foraging patches and temporal variation in prey availability.

## Acknowledgments

We gratefully acknowledge Fumiya Hamai, Yuna Yabuta, Yuuka Mizuguchi, Yasumasa Kitamura, Kana Matsuoka, Shoko Fujitani, Kiho Fujii, Akito Nomi, Yuto Wakasa, Ryota Yao, Mikuto Hori, Tsubasa Sakamoto, for their support in fieldwork and its analysis. Also, we thank Yuichi Mizutani for his help in improving our manuscript.

## Supporting Information

**S1 Dataset**. Times of pond entry, pond exit, and prey attacks by bats.

**S2 Table**. Prey-attack rates calculated by the GLMM for each day.

**S3 Table**. The three parameters calculated using experiment and simulation data.

## References

1. MacArthur RH, Pianka ER. On Optimal Use of a Patchy Environment. The American Naturalist. 1966;100(916):603–9. doi: 10.1086/282454.

2. Charnov EL. Optimal foraging, the marginal value theorem. 1976.

3. Couzin ID. Collective cognition in animal groups. Trends in Cognitive Sciences. 2009;13(1):36–43. doi: 10.1016/j.tics.2008.10.002.

4. Katz Y, Tunstrøm K, Ioannou CC, Huepe C, Couzin ID. Inferring the structure and dynamics of interactions in schooling fish. Proceedings of the National Academy of Sciences. 2011;108(46):18720–5. doi: doi:10.1073/pnas.1107583108.

5. Czaczkes TJ, Grüter C, Ratnieks FL. Trail pheromones: an integrative view of their role in social insect colony organization. Annu Rev Entomol. 2015;60:581–99. Epub 20141024. doi: 10.1146/annurev-ento-010814-020627. PubMed PMID: 25386724.

6. Doering GN, Prebus MM, Suresh S, Greer JN, Bowden R, Linksvayer TA. Emergent collective behavior evolves more rapidly than individual behavior among acorn ant species. Proceedings of the National Academy of Sciences. 2024;121(48):e2420078121. doi: doi:10.1073/pnas.2420078121.

7. Cavagna A, Cimarelli A, Giardina I, Parisi G, Santagati R, Stefanini F, et al. Scale-free correlations in starling flocks. Proceedings of the National Academy of Sciences. 2010;107(26):11865–70. doi: doi:10.1073/pnas.1005766107.

8. Bialek W, Cavagna A, Giardina I, Mora T, Silvestri E, Viale M, et al. Statistical mechanics for natural flocks of birds. Proceedings of the National Academy of Sciences. 2012;109(13):4786–91. doi: doi:10.1073/pnas.1118633109.

9. Kenward R. Hawks and doves: factors affecting success and selection in goshawk attacks on woodpigeons. The Journal of Animal Ecology. 1978:449–60.

10. Sutton GJ, Hoskins AJ, Arnould JPY. Benefits of Group Foraging Depend on Prey Type in a Small Marine Predator, the Little Penguin. PLOS ONE. 2015;10(12):e0144297. doi: 10.1371/journal.pone.0144297.

11. Clark CW, Mangel M. Foraging and flocking strategies: information in an uncertain environment. The American Naturalist. 1984;123(5):626–41.

12. Egert-Berg K, Hurme ER, Greif S, Goldstein A, Harten L, Herrera Montalvo LG, et al. Resource Ephemerality Drives Social Foraging in Bats. Current Biology. 2018;28(22):3667–73. doi: 10.1016/j.cub.2018.09.064.

13. Ward P, Zahavi A. The importance of certain assemblages of birds as “information-centres” for food-finding. Ibis. 1973;115(4):517–34.

14. Bijleveld AI, van Gils JA, Jouta J, Piersma T. Benefits of foraging in small groups: An experimental study on public information use in red knots Calidris canutus. Behavioural Processes. 2015;117:74–81. doi: 10.1016/j.beproc.2014.09.003.

15. Corcoran AJ, Conner WE. Bats jamming bats: Food competition through sonar interference. Science. 2014;346(6210):745–7. doi: doi:10.1126/science.1259512.

16. Beauchamp G. Social predation: how group living benefits predators and prey: Elsevier; 2013.

17. Sansom A, Cresswell W, Minderman J, Lind J. Vigilance benefits and competition costs in groups: do individual redshanks gain an overall foraging benefit? Animal Behaviour. 2008;75(6):1869–75. doi: 10.1016/j.anbehav.2007.11.005.

18. Bijleveld AI, Folmer EO, Piersma T. Experimental evidence for cryptic interference among socially foraging shorebirds. Behavioral Ecology. 2012;23(4):806–14. doi: 10.1093/beheco/ars034.

19. Griffin DR. Listening in the dark: the acoustic orientation of bats and men. New Haven: Yale University Press; 1958.

20. Simmons JA, Fenton MB, O’Farrell MJ. Echolocation and pursuit of prey by bats. Science. 1979;203(4375):16–21. PubMed PMID: 758674.

21. Surlykke A, Moss CF. Echolocation behavior of big brown bats, Eptesicus fuscus, in the field and the laboratory. J Acoust Soc Am. 2000;108(5 Pt. 1):2419–29. PubMed PMID: 11108382.

22. Fujioka E, Aihara I, Sumiya M, Aihara K, Hiryu S. Echolocating bats use future-target information for optimal foraging. Proc Natl Acad Sci USA. 2016;113(17):4848–52. doi: 10.1073/pnas.1515091113.

23. Luo J-h, Ou W, Liu Y, Wang J, Wang L, Feng J. Plasticity in echolocation calls of Myotis macrodactylus (Chiroptera: Vespertilionidae): implications for acoustic identification. Acta Theriologica. 2012;57(2):137–43.

24. Mizuguchi Y, Fujioka E, Heim O, Fukui D, Hiryu S. Discriminating predation attempt outcomes during natural foraging using the post-buzz pause in the Japanese large-footed bat, Myotis macrodactylus. Journal of Experimental Biology. 2022;225(7). doi: 10.1242/jeb.243402.

25. Balcombe JP, Fenton MB. Eavesdropping by bats: the influence of echolocation call design and foraging strategy. Ethology. 1988;79(2):158–66.

26. Gillam E. Eavesdropping by bats on the feeding buzzes of conspecifics. Canadian Journal of Zoology. 2007;85:795–801. doi: 10.1139/Z07-060.

27. Dechmann DKN, Heucke SL, Giuggioli L, Safi K, Voigt CC, Wikelski M. Experimental evidence for group hunting via eavesdropping in echolocating bats. Proceedings of the Royal Society B: Biological Sciences. 2009;276(1668):2721–8. doi: 10.1098/rspb.2009.0473.

28. Roeleke M, Schlägel UE, Gallagher C, Pufelski J, Blohm T, Nathan R, et al. Insectivorous bats form mobile sensory networks to optimize prey localization: The case of the common noctule bat. Proceedings of the National Academy of Sciences. 2022;119(33):e2203663119. doi: doi:10.1073/pnas.2203663119.

29. Krivoruchko K, Koblitz JC, Goldshtein A, Biljman K, Guillén-Servent A, Yovel Y. A social foraging trade-off in echolocating bats reveals that they benefit from some conspecifics but are impaired when many are around. Proceedings of the National Academy of Sciences. 2024;121(30):e2321724121. doi: doi:10.1073/pnas.2321724121.

30. Fukui D, Agetsuma N, Hill DA. Acoustic Identification of Eight Species of Bat (Mammalia: Chiroptera) Inhabiting Forests of Southern Hokkaido, Japan: Potential for Conservation Monitoring. Zoological Science. 2004;21(9):947–55. doi: 10.2108/zsj.21.947.

31. Eberhard JR, Ewald PW. Food availability, intrusion pressure and territory size: an experimental study of Anna’s hummingbirds (Calypte anna). Behavioral Ecology and Sociobiology. 1994;34(1):11–8. doi: 10.1007/BF00175453.

32. Götze S, Denzinger A, Schnitzler H-U. High frequency social calls indicate food source defense in foraging Common pipistrelle bats. Scientific Reports. 2020;10(1). doi: 10.1038/s41598-020-62743-z.

33. Hamilton WD. The genetical evolution of social behaviour. I. Journal of Theoretical Biology. 1964;7(1):1–16. doi: 10.1016/0022-5193(64)90038-4.

34. Tibbetts EA, Pardo-Sanchez J, Weise C. The establishment and maintenance of dominance hierarchies. Philosophical Transactions of the Royal Society B: Biological Sciences. 2022;377(1845). doi: 10.1098/rstb.2020.0450.

35. Wilkinson GS. Reciprocal food sharing in the vampire bat. Nature. 1984;308(5955):181–4.

36. Oro D. LIVING IN A GHETTO WITHIN A LOCAL POPULATION: AN EMPIRICAL EXAMPLE OF AN IDEAL DESPOTIC DISTRIBUTION. Ecology. 2008;89(3):838–46. doi: 10.1890/06-1936.1.

37. Guo D, Luo B, Zhang K, Liu M, Metzner W, Liu Y, et al. Social vocalizations of big-footed myotis (Myotis macrodactylus) during foraging. Integrative Zoology. 2019;14(5):446–59. doi: 10.1111/1749-4877.12367.

38. Guo D, Ding J, Liu H, Zhou L, Feng J, Luo B, et al. Social calls influence the foraging behavior in wild big-footed myotis. Frontiers in Zoology. 2021;18(1). doi: 10.1186/s12983-020-00384-8.

39. Fretwell SD. Populations in a seasonal environment: Princeton University Press; 1972.

40. Encarnacao JA, Becker NI, Ekschmitt K. When do Daubenton’s bats (Myotis daubentonii) fly far for dinner? Canadian Journal of Zoology. 2010;88(12):1192–201.

